# Spatial-CITE-seq: spatially resolved high-plex protein and whole transcriptome co-mapping

**DOI:** 10.1101/2022.04.01.486788

**Authors:** Yang Liu, Marcello DiStasio, Graham Su, Hiromitsu Asashima, Archibald Enninful, Xiaoyu Qin, Yanxiang Deng, Pino Bordignon, Marco Cassano, Mary Tomayko, Mina Xu, Stephanie Halene, Joseph E. Craft, David Hafler, Rong Fan

## Abstract

We present spatial-CITE-seq for high-plex protein and whole transcriptome co-mapping, which was firstly demonstrated for profiling 189 proteins and transcriptome in multiple mouse tissue types. It was then applied to human tissues to measure 273 proteins and transcriptome that revealed spatially distinct germinal center reaction in tonsil and early immune activation in skin at the COVID-19 mRNA vaccine injection site. Spatial-CITE-seq may find a range of applications in biomedical research.

## Main Text

Spatially resolved transcriptome sequencing is transforming the study of cell differentiation and tissue development^1-3^, but does not yet incorporate protein expression profiling. Previously, we developed microfluidic deterministic barcoding in tissue (DBiT) for co-mapping of whole transcriptome and a panel of 22 proteins at the cellular level (∼10µm pixel size) using antibody-derived DNA tags (ADTs)^4^ to convert the detection of proteins to the sequencing of corresponding DNA tags^5, 6^. Array-based spatial transcriptome was also expanded to multi-omics, namely SM-Omics^7^, which demonstrated the mapping of 6 proteins and whole transcriptome with 100 µm spot size. Very recently, Landau and co-authors further implemented spatial multi-omics on the 10X Visium platform with 55µm spot size and a panel of 21 protein markers^8^. However, it remains unclear how large a panel of proteins can be simultaneously mapped and what difference can be obtained if ultra-high-plex (>100) protein mapping was realized.

Herein, we report on spatial-CITE-seq: spatial co-indexing of transcriptomes and epitopes for multi-omics mapping by next generation sequencing (NGS), which uses a cocktail of ∼200-300 ADTs to stain a tissue slide followed by deterministic in tissue barcoding of both DNA tags and mRNAs for spatially resolved high-plex protein and transcriptome co-profiling (**Figure 1a**). Each ADT contains a poly-A tail, a unique molecular identifier (UMI), and a specific DNA sequence unique to the corresponding antibody (**Figure S1**). A large panel of ADTs were combined in a cocktail and applied to a paraformaldehyde (PFA)-fixed tissue section (∼7µm in thickness). Next, a microfluidic chip was used to introduce to the tissue surface a panel of DNA row barcodes A1- 50, each of which contains an oligo-dT sequence that binds to the poly-A tail of ADTs or mRNAs, followed by in tissue reverse transcription. Then, a panel of DNA column barcodes B1-50 were flowed over the tissue surface in a perpendicular direction using a different microfluidic chip and ligated in situ to create a 2D grid of tissue pixels, each containing a unique spatial address code AiBj (i=1-50, j=1-50) to co-index all protein epitopes and transcriptome. Finally, barcoded cDNAs were recovered, purified, and PCR amplified to prepare two NGS libraries for paired-end sequencing of ADTs and mRNAs, respectively, for computational reconstruction of spatial protein or gene expression map.

**Figure 1.**
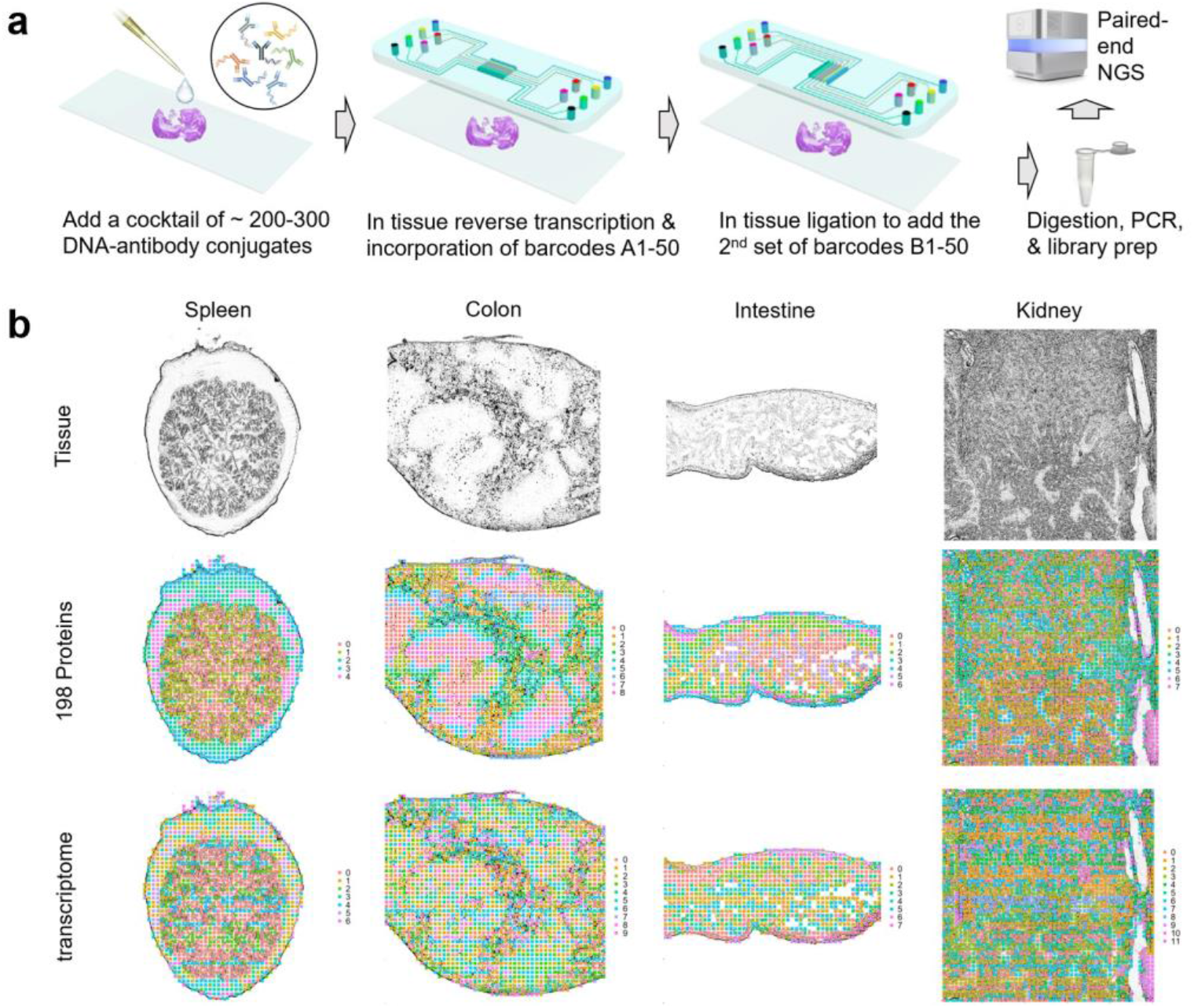
Spatial-CITE-seq workflow design and application to diverse mouse tissue types for co-mapping of 189 proteins and whole transcriptome. (a) Scheme of spatial-CITE-seq. A cocktail of antibody-derived DNA tags (ADTs) is applied to a PFA-fixed tissue section to label a panel of ∼200-300 protein markers in situ. Next, a set of DNA barcodes A1-A50 are flowed over the tissue surface in a spatially defined manner via parallel microchannels and reverse transcription is carried out inside each channel for in-tissue synthesis of cDNAs complementary to endogenous mRNAs and introduced ADTs. Then, a set of DNA barcodes B1-B50 is introduced using another microfluidic device with microchannels perpendicular to the first flow direction and subsequently ligated to barcodes A1-A50, creating a 2D grid of tissue pixels, each of which has a unique spatial address code AB. Finally, barcoded cDNA is collected, purified, amplified, and prepared for paired end NGS sequencing. (b) Spatially resolved 189-plex protein and whole transcriptome co-mapping of mouse spleen, colon, intestine, and kidney tissue with 20µm pixel size. Upper row: brightfield optical images of the tissue sections. Middle row: unsupervised clustering of all pixels based on all 189 protein markers only and projection onto the tissue images. Lower row: unsupervised clustering of whole transcriptome of all pixels and projection to the tissue images. Colors correspond to different proteomic or transcriptomic clusters indicated on the right side of each panel.

It was first demonstrated for spatial mapping of 189 proteins and genome-wide gene expression in multiple mouse tissue types including spleen, colon, intestine, kidney, etc. The mouse ADT panel (**Table S5**) includes the markers for canonical cell types and immune cell function. The total number of proteins detect is approaching ∼190, indictive of high sensitivity to detect even non-specific background noises. In the mouse spleen sample, the average protein count per pixel (25 µm) is 118 and the protein UMI account per pixel is 885 (**see Table S1**). Low UMI count pixels are localized in the low cell density capsule region (see **Figure S2**). Uniquely, unlike our previous work that mapped much smaller number of proteins and did not perform well tissue region clustering analysis using the protein profiles alone, this high-plex protein panel allowed for unbiased clustering of all tissue pixels into spatially distinct clusters. Spatial protein profiles in the spleen sample resulted in 5 major clusters (**Figure 1b**). Clusters 0 and 1 separates red and white pulps. Cluster 2 indicates microvascular tissue. Clusters 3 and 4 are enriched in spatially distinct regions of the capsule. Spatial transcriptome data from the same tissue section is of high quality (average gene count and UMI count per pixel: 1166 and 1972) (**Table S1**). Transcriptome clustering analysis identified 7 clusters that also resolved red and white pulps in concordance with spatial high-plex protein clustering. Mouse colon, intestine and kidney tissues were also analyzed, and the resultant major clusters correlated with anatomic regions (**Figure 1b**).

We further conducted spatial co-mapping of 273 human protein markers (**Table S5**) and whole transcriptome in human secondary lymphoid (tonsil) tissue over a 2mmx2mm region of interest (indicated by a dashed box in **Figure 2a**). Average protein count per pixel is 239 with the average UMI count 4309 (**Figure 2b, Table S1**). Clustering of spatial protein profiles alone identified 7 major clusters (**Figure 2c**) and the corresponding spatial distribution showed highly distinct features (**Figure 2d**). Spatial transcriptome obtained in this experiment gave rise to 8 major clusters (**Figure 2e**) and their spatial distribution (**Figure f**) correlated well with spatial protein clusters but appeared to be more noisy and less precise. Differential protein expression analysis (**Figure 2g**) allowed for identification of major cell types in each cluster. Overlay of tissue image and spatial protein cluster map (**Figure 2h**) showed a strong correlation between anatomic features and tissue/cell types. Cluster 0 corresponds to the crypt epithelia. Clusters 2 and 5 are the germinal center (GC) light and dark zones. Cluster 1 indicates specific T-cell zones. Clusters 3 and 4 are localized in extrafollicular regions. Cluster 6 contains peripheral blood cells in vasculature. We further visualized individual proteins one by one. For example, CD19, a marker for B cells, is enriched in follicles^9^. CD21 or complement receptor 2 (CR2)^10^, present on all mature B cells as well as follicular dendritic cells (DCs), is highly expressed in the whole follicles. CD23, previously found on mature B cells, activated macrophages, eosinophils, follicular dendritic cells, and platelets, is restricted to the apical region of the GC light zone^11^. We further examined the functional proteins such as immunoglobulins associated with B cell differentiation and maturation (**Figure 2j**). IgM expression is restricted to GC B cells. Once they further mature, these B cells start to produce IgG and migrate out of follicles. IgD is produced mainly by naïve B cells that just exit from the blood stream. CD90 (Thy-1) is associated with a wide range of cell types but completely absent in GCs. Notch3 is found in squamous epithelial cells. Mac2/Galectin3 is highly enriched in the crypt zone. We also examined T cell marker CD3 that identified all major T-cell zones as well as CD4 for helper T cells and CD45A for naïve or stem cell-like T cells (**Figure 2l**). CD32 is an Fc receptor that regulates B cell activation^12^ and was found mainly outside GCs. CD9 is expressed in tonsillar B cells in both follicles and crypts. CD171, a neuronal cell adhesion molecule implicated in neurite outgrowth, myelination and neuronal differentiation, is found to be highly restricted in the dark zone. This has not been reported previously and warrants further investigation (**Figure 2m**). Selected proteins were validated by immunofluroscence staining performed on COMET™ (**Figure S3**)^13^. In addition, we also demonstrated the applicability of spatial-CITE-seq to other human tissues including spleen and thymus (see **Figure S4**).

**Figure 2.**
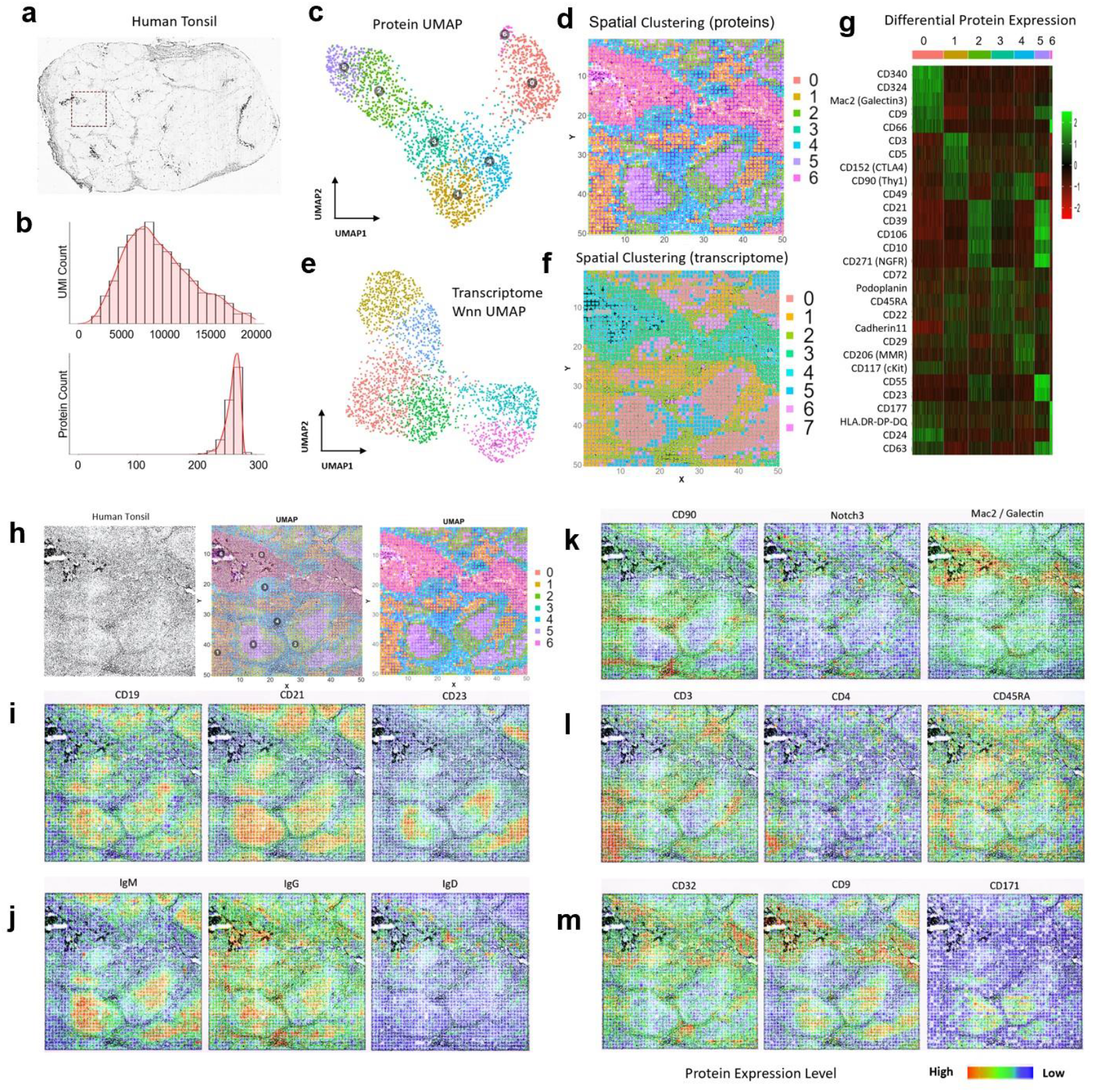
Spatial co-mapping of 273 protein markers and whole transcriptome in human tonsil revealed spatially distinct germinal center (GC) reaction and extrafollicular activities. (a) Image of a human tonsil tissue section. The region mapped by spatial-CITE-seq is indicated by a dashed box. (b) per-pixel UMI count and protein count histograms. (c) UMAP plot of the clustering analysis of all pixels based on 273 proteins only. (d) Spatial distribution of the clusters (0-6) indicated by the same colors as in (c). (e) UMAP plot of the clustering analysis of all pixels based on the mRNA transcriptome. (f) Spatial distribution of the transcriptomic clusters (0-7) indicated by the same colors as in (e). Pixel size: 20µm (g) Differentially expressed proteins in the clusters shown in (c, d). (h) Tissue mage of the mapped region (left), spatial proteomic clusters (right), and the overlay (middle). (i) Individual surface protein markers related to B cells and follicular DCs. (j) Functional protein markers such as immunoglobulins showing spatially distinct distribution of GC B cells (IgM), matured B cells (IgG), and naïve B cells (IgD), in agreement with B cell maturation, class switch, and migration. (k) Individual protein markers enriched in the extracellular region (CD90, Notch3) and crypt (Mac2). (l) Individual T cell protein markers CD3, CD4, and CD45RA showing T-cell zones and subtypes. (m) Individual protein markers CD32, CD9, and CD171. CD32 identified a range of immune cells including platelets, neutrophils, macrophages, and dendritic cells (DCs) trafficking from vasculature. CD9 identified plasma cell precursors in germinal centers (GCs) and crypt. CD171, a neural cell adhesion molecule, is found highly distinct in the GC dark zone. Color key: protein expression from high to low.

Finally, spatial-CITE-seq was used to map early immune cell activation in a skin biopsy tissue collected from the COVID-19 mRNA vaccine injection site. The tissue section is comprised of collagen-rich region with low cell density and a vascular granule region with high cellularity (**Figure 3a**). We evaluated the data quality for both transcriptome and proteins (**Figure S5**). Spatial map of gene count correlates with cell density and the high cell density region resulted in 411 genes per pixel (**Figure 3b**). However, unsupervised clustering identified spatially distinct clusters even in the low cell density regions (**Figure 3c**). Spatial map of protein count is less variable across the tissue section and up to ∼270 proteins could be detected in the low-density region (**Figure 3e**). Clustering of spatial protein profiles gave rise to 10 clusters (**Figure 3d**) and the corresponding spatial distribution (**Figure 3f**) was highly distinct in strong agreement with the spatial transcriptome clusters.

**Figure 3.**
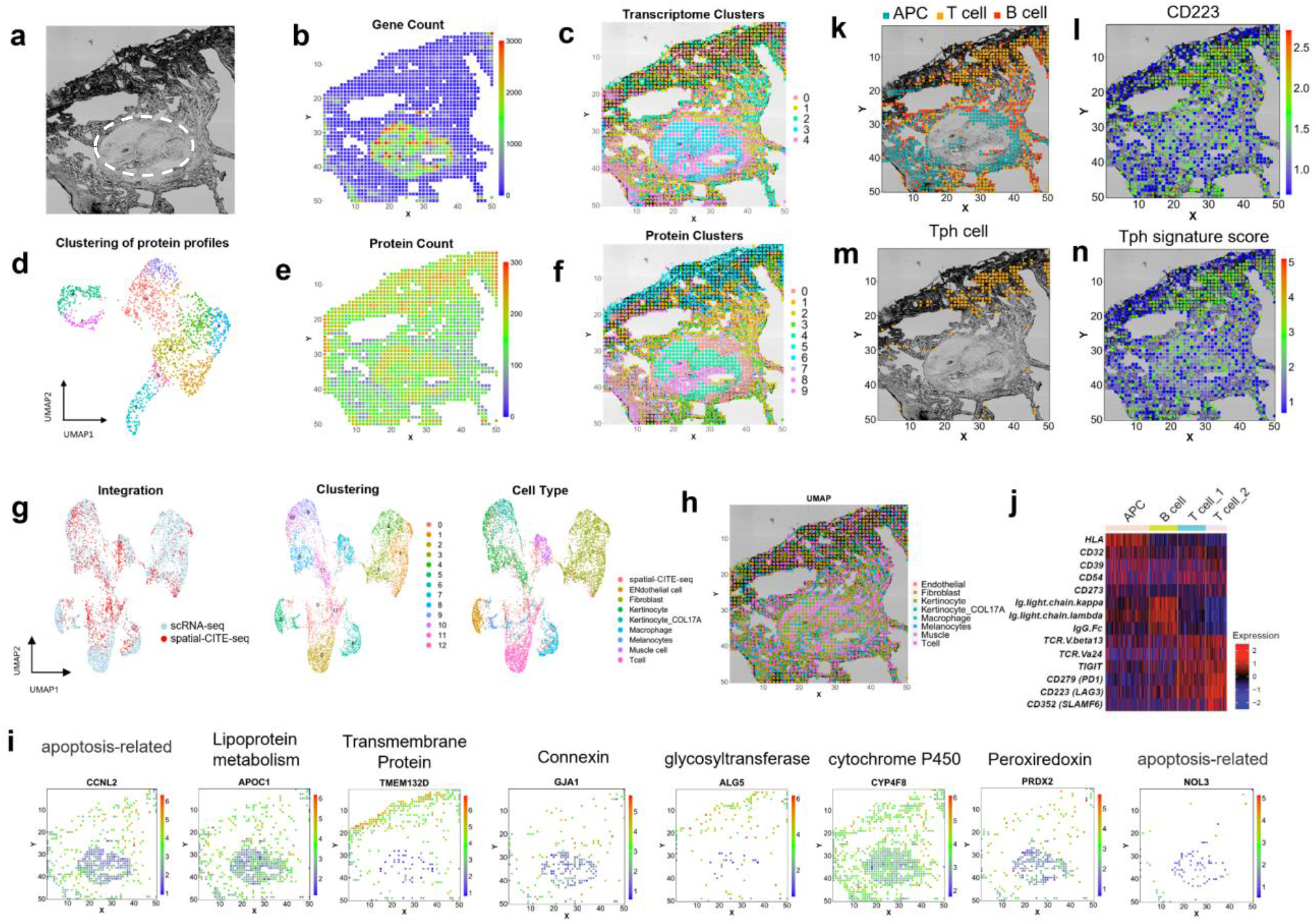
Integrated spatial and single-cell profiling of a human skin biopsy tissue at the site of COVID19 mRNA vaccination injection revealed localized peripheral T cell activation. (a) Brightfield image of skin section in the mapped region. A pilosebaceous unit is indicated by the dashed region (b) Gene count spatial map. (c) Spatial clustering of all pixels based on whole transcriptome. Despite low gene count in the low cell density regions of dermal collagen, the clustering analysis revealed spatially distinct zones based on transcriptomic profiles. (d) UMAP clustering of all 273 proteins. (e) Protein count distribution. (f) Spatial clustering of all pixels based on 273 proteins only, which is in high concordance with spatial clusters identified by spatial transcriptome co-mapped on the same tissue section. (g) Integrated analysis of single-cell and spatial transcriptome. Left: the transcriptomes of spatial tissue pixels (red) conform to the clusters identified by joint analysis with single-cell RNA-seq (blue). Middle: unsupervised clustering of the combined transcriptome dataset. Right: cell type annotation. (i) Visualization of select genes associated with different gene oncology functions via integrated analysis and transfer learning. (j) Differential protein expression in different cell types (APC, B cell, and two subtypes of T cells). (k) Spatial distribution of APC, T, and B cells. (l) Expression of CD223 (LAG3) protein, a functional marker of activated T cells and other immune cell subsets. (m) Identification of a highly localized population of peripheral helper T cells (Tph) at the vaccine injection site. (n) Spatial distribution of Tph gene score correlates with the cell localization. Pixel size: 20µm.

Single-cell RNA sequencing (scRNA-seq) was conducted with the same skin biopsy tissue specimen (**Figure S6**). It was combined with spatial transcriptomes to perform clustering that gave rise to 13 major clusters and the major cell types were identified based on gene oncology (**Figure 3g**). Label transfer of cell types from scRNA-seq to spatial tissue pixels allowed for visualization of the distribution of different types (**Figure 3h**). We can also visualize the expression of individual genes (**Figure 3i**). For example, CCNL2 and NOL3, which are apoptosis related genes were expressed in the vascular region; APOC1(responsible for lipoprotein metabolism), GJA1(Connexin protein encoding) and PRDX2 (Peroxiredoxin encoding), were expressed mainly in the vascular. Transmembrane protein TMEM132D and glycosyltransferase ALG5 were both expressed in the dermis region. CYP4F8, encoding CYP450 protein were shown in most skin regions. The whole transcriptome sequencing could identify the cell types in general but were not specific enough here to show the different populations of T cells. Next, we focused on several immune cell types including antigen presenting cells (APCs), B cells, and two subsets of T cells, as indicated by differentially expressed proteins (**Figure 3j**). APCs and T cells are localized in spatially distinct regions while B cells are distributed throughout the tissue (**Figure 3k**), Specifically, T cell subset 2 express a set of markers including lymphocyte activation gene 3 (LAG3)^14^, associated with peripheral helper T cell population (Tph)^15^ (**Figure 3m**) as definitely by Tph signature score defined by expression levels of LAG3, PD-1 and CXCR6 (**Figure 3n**). Tph cells are implicated in local T cell activation in response to vaccination. Thus, through integration of spatial high-plex protein and transcriptome mapping with single-cell RNA-seq data from same skin biopsy tissue, we identified major skin and immune cell types and a subset of peripheral helper T cells highly enriched at the injection site which may contribute to the local immune activation that initiates systemic vaccine response.

Latest advances in imaging-based protein mapping such as imaging mass cytometry (IMC)^16^ or multiplex immunofluorescence (i.e., CODEX^17^, CyCIF^18, 19^, and seqIF^20^) has realized 25-60-plex protein mapping and transformed spatial protein biomarker research. Our work utilized spatial barcoding and high-throughput sequencing for the mapping of ∼200-300 proteins, representing the highest multiplexing to date for spatial protein profiling despite the lack of subcellular resolution. It could be expanded to >1000-plex protein mapping given that only ∼10% of the sequencing lane was used for the ADT library. We noticed a competition between ADTs and mRNAs for in tissue reverse transcription and lower efficiency to detect transcripts compared to to single modality spatial transcriptome sequencing. This requires future optimization such as ADT concentration and enzymatic reaction conditions. Current protein panel is largely comprised of surface epitopes and yet to be further expanded to intracellular proteins or extracellular matrix proteins to investigate a wide range of protein signaling and function. In short, spatial-CITE-seq incorporates ∼200-300 protein markers and offers significant enhancement in the capabilities of tissue mapping, with applications to unmet needs in a wide range of fields including cancer, immunology, infectious disease, and anatomic pathology.

## METHODS

### Microfluidic device design and fabrication

We designed the photomask using Autodesk AutoCAD 2021 and had the chrome mask printed by the company Front Range Photomasks (Lake Havasu City, AZ) with high resolution (2 µm). The chrome mask was cleaned extensively with acetone and air-dried before use. PDMS mold (25 µm channel width) was fabricated in a cleanroom using Photoresist SU-8 2025 (Kayaku Advanced Materials, Inc) following standard procedures including, spin coating, soft baking, laser exposure, post-exposure baking, development, and hard baking. The mold thickness was measured using Zygo 3D Optical Profiler to be ∼25 µm. The mold was placed in a plastic petri dish and the PDMS mixture (Part A: part B = 10:1, GE RTV) was poured in. The petri dish was placed into a vacuum chamber and degassed for ∼30 mins and then placed into a 70°C oven and incubated for >2 hours or overnight. The cured PDMS slab was cut into a similar size as a 1×3 inches glass slide and stored at room temperature until use. The barcoding flow clamps, and lysis clamps were fabricated through laser-cutting an acrylic plastic plate. After each DBiT-seq experiment, the PDMS chip can be reused by cleaning with 30 mins sonication in 1M NaOH solution, 2 hours soaking in DI water, 10 mins sonication in isopropanol, and air dry at room temperature.

### Microscope setup

The tissue image and two flow channel/tissue images were scanned with the Invitrogen EVOS M7000 imaging system using a 10x objective. Images were taken with mono-color mode and stitched with “More Overlap” settings. The stitched images were saved into TIFF format and later aligned with spatial transcriptome and proteome data.

### DNA oligos and antibody-derived DNA tags

DNA oligos used were all synthesized by Integrated DNA Technologies with HPLC purification. All DNA oligos received were dissolved in RNase-free water at a 100 µM concentration and stored at -20 °C until use. All the DNA oligos used were listed in this file (Table S2). The barcode A and B oligos were listed in Table S3. Barcode A contains three functional regions: a poly-T region, a spatial barcode region, and a ligation linker region. Poly-T region hybrids with poly-A tail of mRNA and serves as the RT primer. The spatial barcode defines the row locations, and the ligation linker region was to be ligated with barcode B. Barcode B includes four functional regions: one ligation linker region, a spatial barcode region, a UMI region and a PCR primer region. The ligation linker region was to be ligated to barcode A. The spatial barcode region shows the column locations. Barcode B was also functionalized with 5’ biotin.

Antibody-derived DNA tags for membrane proteins were purchased from Biolegend and listed in Table S5. Three antibody cocktail products are 273 antibodies cocktail for humans with 9 isotype control antibodies (Cat No. 99502), and 189 antibodies cocktail for mice with 9 isotype control antibodies (Cat No. 99833).

### Tissue preparation

OCT embedded mouse spleen (Mouse CD1 Spleen Frozen Sections, MF-701), colon (Mouse CD1 Colon Frozen Sections, MF-311), intestine (CD1 Intestine, Jejunum Frozen Sections, MF- 308), kidney (Mouse CD1 Kidney Frozen Sections, MF-901) sections were purchased from Zyagen Inc (San Diego, CA), and stored at -80°C until use. In a typical protocol, OCT tissue blocks were sectioned into 10 µm thickness sections and placed in the center of poly-L-lysine slides (Electron Microscopy Sciences, 63478-AS), and shipped with dry ice. The human tonsil sections (Human Tonsil Frozen Sections, HF-707) were also purchased from Zyagen (San Diego, CA). Human skin samples were obtained from the Yale neurology department and sectioned into a 10- µm thickness.

### Spatial-CITE-seq profiling of tissue

OCT embedded tissue sections stored in a -80 °C freezer were left on the working bench for 10 minutes. Sections were then fixed with 4% formaldehyde for 20 minutes and washed three times with 1x PBS with 0.05U/μL RNase Inhibitor (Enzymatics, 40 U/μL). The tissue was then permeabilized with 0.5% Triton X-100 in 1x PBS for another 20 minutes before washing three times with 1x PBS. The sections were quickly dipped in RNase free water and dried with air. We then covered the tissue using 1x blocking buffer with 0.05U/μL RNase Inhibitor (Enzymatics, 40 U/μL) and incubated at 4 °C for 10 minutes. After washing three times with 1xPBS buffer, ADT cocktails (diluted 20 times from original stock) from Biolegend were added onto the tissue and incubated for 30 minutes at 4 °C. The ADT cocktail was removed by washing three times with 1X PBS and the slide was dipped in water briefly to remove any remaining salts. A whole tissue image scan was performed with EVOS microscope using a 10x objective.

In tissue, reverse transcription was conducted by flowing reverse transcription reagents into each of the 50 channels. We prepared the Reverse Transcription mix by adding sequentially 50 μL of 5X RT Buffer (ThermoFisher), 7.8 μL of RNase-free water, 1.6 μL RNase Inhibitor (Enzymatics Inc), 3.2 μL μL SuperaseIn RNase Inhibitor (Ambion), 12.5 μL of 10 mM dNTPs each (ThermoFisher), 25 μL of Maxima H Minus Reverse Transcriptase (ThermoFisher), and 100 μL 0.5X PBS-RI (0.5X PBS + 1% Rnase Inhibitor from Enzymatics) into a 1.5 mL tube. The mix was enough for a DBiT-seq chip with 50 channels and was further mixed with individual Barcode A (25 μM in water) with a 4:1 volume ratio. The first PDMS chip was then placed on top of the tissue section and customized plastic clamps were applied to the chip to seal tightly the PDMS chip with the tissue. The slide was imaged again with EVOS microscope to record the locations of the channels. A total volume of 5 µL of Reverse Transcription mix and Barcode A was loaded into each inlet well on the first PDMS chip. After loading and carefully removing air bubbles inside each well, a vacuum adapter made with acrylic plastic was placed on the outlet wells of the chip, and solutions were then vacuumed through the 50 channels. After 2 minutes, the vacuum was turned off and the chip was placed into a wet box and incubated first at room temperature for 30 minutes and then 90 minutes at 42 °C. When the RT reaction was completed, the channels were flushed with 1X NEB buffer 3.1 with 1% RNase Inhibitor (Enzymatics) for 5 minutes. After removing the 1^st^ PDMS chip, the tissue was dipped in RNase-free water and kept dry at 4 °C until the next step.

In tissue ligation was performed in the 2^nd^ PDMS chip which has 50 channels with orthogonal direction. The barcode B and ligation linker mix was first prepared by mixing barcode B (100 µM in water), 10 µL ligation linker oligo (100 µM in water), and 20 µL annealing buffer (10 mM Tris, pH 7.5–8.0, 50 mM NaCl, 1 mM EDTA) in a PCR tube and then heated to 90–95 °C for 3–5 minutes before cooling to room temperature on the workbench. The mix was stored at 4 °C for short-term use or at -20 °C for long-term storage.

The ligation mix was prepared by adding into a 1.5 mL Eppendorf tube 68 µL of RNase-free water, 29 μL 10X T4 Ligase buffer (NEB), 11 μL T4 DNA Ligase (400 U/μL, NEB), 2 μL RNase inhibitor (40 U/μL, Enzymatics), 0.7 μL SuperaseIn RNase Inhibitor (20 U/μL, Ambion), 5.4 μL of 5% Triton- X100, and 116 µL 1X NEB buffer 3.1 with 1% RNase Inhibitor (40 U/μL, Enzymatics). 4 µL of ligation mix was mixed with 1 µL Barcode B (25 µm, with ligation linker) in a 96 well plate. The 2^nd^ PDMS chip was attached to the section and clumped together with an acrylic clump. The chip was scanned with the EVOS microscope to record the spatial locations of channels. 5 µL of the above mixture was loaded into the inlet wells of the PDMS chip and vacuumed through each channel. The chip was transferred to a 37 °C oven and incubated for 30 minutes. The remaining solution in the inlets wells was removed and wash buffer (1X PBS with 0.1% Triton-X 100) was loaded and vacuumed through the channels continuously for 5 minutes. The PDMS chip was peeled off and the tissue was dipped in water and dried with air.

The whole tissue section was digested by proteinase K to release the cDNAs. We prepare the lysis buffer by mixing 50 μL 1x PBS, 50 μL of 2X lysis buffer (20 mM Tris pH 8.0, 400 mM NaCl, 100 mM EDTA, and 4.4% SDS) and 10 μL of proteinase K solution (20mg/mL). A PDMS reservoir was placed on top of the ROI and the lysis mix was added. The reservoir was then clamped tightly with the slide to avoid any leakage and was sealed with parafilm. The tissue was lysed in a 55 °C oven for 2 hours, and the lysis was collected and kept in a -80 °C freezer until use.

cDNA extraction from the tissue lysate was performed in two steps. In the first step, all DNA was extracted from the lysate using the DNA purification kit (ZYMO Research, Cat. ZD4014). We followed recommended protocols using a 5:1 ratio for the DNA binding buffer and lysate. In the second step, biotinylated cDNAs were captured with streptavidin beads (Dynabeads MyOne Streptavidin C1, Invitrogen). Before use, the beads were washed three times with 1x B&W buffer with 0.05% Tween-20 and dispersed into 100 µL of 2X B&W buffer. The beads were added into the purified cDNA with a 1:1 volume ratio and incubated with mild rotation at room temperature for 1 hour. Beads were cleaned twice with 1X B&W buffer and once using 1X Tris buffer with 0.1% Tween-20.

To add a 2^nd^ PCR handle to the cDNA strands, template switch was performed. We prepared the template switch reagents with standard protocol, using 44 µL 5X RT buffer, 44 µL Ficoll PM-400 solutions, 22 µL dNTPs, 5.5 µL RNase inhibitor, 11 µL Maxima H Minus Reverse Transcriptase,

5.5 µL of template switch oligo, and 88 µL of H2O. The beads were resuspended into the mix and the reaction was performed at room temperature for 30 minutes and then 1.5 hours at 42 °C with rotation. After template switch, the beads were cleaned once with 1X PBST (0.1% Tween 20) and once with water.

We prepared the 220 µL of PCR mix with 110 µL Kapa HiFi HotStart Master mix, 8 µL primer 1 (10 µM), 8 µL primer 2 (10 µM), 0.5 µL primer 2-citeseq (1 µM), and 91.9 µL H2O. The cleaned Dynabeads were redispersed in this PCR mix and the solution was split into four PCR tubes with 55 µL each. PCR was performed by first incubating at 95 °C for 3 minutes, then running 20 cycles at 98 °C for 20 seconds, 65 °C for 45 seconds, and 72 °C for 3 minutes. To separate the cDNAs derived from RNA and cDNAs derived from ADT, we did the purification using 0.6x SPRI beads following standard protocol. Specifically, we added 120 µL of SPRI beads to 200 µL of PCR product solution and incubated for 5 minutes. The supernatant containing the ADT cDNAs was collected in a 1.5 mL Eppendorf tube. The remaining beads were cleaned with 85% ethanol for 0.5 minute and then eluted with RNase-free water for 5 minutes. The cDNAs derived from mRNA were then quantified with Qubit and bioanalyzer. For the supernatant, we added another 1.4X SPRI beads and incubated them for 10 minutes. The beads were cleaned once with 80% ethanol and redispersed in 50 µL water. We did another 2X SPRI purification by adding 100 µL SPRI beads and incubated for 10 minutes. After washing twice with 80% ethanol, we collected the cDNAs derived from ADTs by eluting them with 50 µL RNase-free water.

The sequencing library of the two types of cDNA products was built separately. For cDNAs derived from mRNA, 1 ng of the cDNA was used, and the library was built using the Nextera XT Library Prep Kit (Illumina, FC-131-1024) using customized index strands and purified with 0.6x SPRI beads. For ADT cDNAs, the library was built with PCR. In a PCR tube, 45 µL of ADT cDNA solution, 50 µL of 2x KAPA Hifi PCR master mix, 2.5 µL customized i7 index (10 µM), and 2.5 µL of P5 index (N501-citeseq, 10 µM) were mixed. PCR was performed at 95 °C for 3 minutes, then cycled at 95 °C for 20 seconds, 60 °C for 30 seconds and 72 °C for 20 seconds for totally 6 cycles, and the reaction was finished with incubation at 72 °C for 5 minutes. The product was purified with 1.6X SPRI beads and then quantified with QuBit and BioAnalyzer. The libraries were sequenced with Novaseq 6000 system.

### Single cell RNA-seq for human skin biopsy sample

Skin punch biopsies were placed immediately into MACS Tissue Storage Solution (Miltenyi Biotec, 130-100-008) and processed into single-cell suspensions using the Whole Skin Dissociation Kit (Miltenyi Biotec, Cat No.130-101-540) according to the manufacturer’s recommendation. Briefly, the tissue was placed in the enzyme solution, and incubated in a 37 °C water bath for 3 hours. Thereafter, the tissue cells were dissociated using the MACS Dissociator (Miltenyi Biotec, Cat No. 130-093-235), preprogrammed for skin cell isolation (program h-skin-01). The cells were then resuspended in DMEM, and mononuclear cells were isolated by Ficoll-Paque PLUS (GE Healthcare) gradient centrifugation. Single-cell preparations were loaded into the Chromium Controller (10x Genomics) for emulsion generation, and libraries were prepared using the Chromium Single Cell 5′ Reagent Kit for version 1.1 chemistry per the manufacturer’s protocol. Libraries were sequenced on the NovaSeq 6000 for gene expression and BCR/TCR libraries.

### Data preprocessing

For cDNAs derived from mRNAs, the raw fastq file of Read 2 containing the UMI, barcode A and barcode B regions, were first reformatted into the standard input format required by ST pipeline v1.7.2 ^21^ using customized python script. Using recommended ST pipeline parameters, the read 1 was STAR mapped to either the mouse genome (GRCm38) or the human genome (GRCh38). The gene expression matrix contains the spatial locations (Barcode A x Barcode B) of the genes and gene expression levels.

For cDNAs derived from ADTs, the raw fastq file of Read 2 was reformatted the same way as cDNAs from RNA. Using default settings of CITE-seq-Count 1.4.2^22^, we counted the ADT UMI numbers for each antibody in each spatial location. The protein expression matrix the spatial locations (Barcode A x Barcode B) of the proteins and protein expression levels.

### Clustering and visualization

The clusters of RNA and protein expression matrix was generated using Seurat V3.2^23^. The transcriptome data was normalized using “SCTransform” function. Normalized data were then clustered and UMAP was built with the dimension were set to 30, and cluster resolution was set to 0.5. Protein data were normalized using the centered log ratio (CLR) transformation method in Seurat V3.2. All the heatmap was plotted using ggplot2.

### scRNA-seq and spatial data integration

The cell types of skin biopsy section were annotated through integration analysis using the matched scRNA-seq data as the reference. The two data sets were normalized with the “SCTransform” function in Seurat V3.2, and then integrated into one dataset. After clustering, the spatial pixel data conformed well with the scRNA-seq data, and thus the cell types were assigned based on the scRNA-seq cell type annotation for each cluster (if two cell types presented in one cluster, the major cell types were assigned).

### Fluorescent staining of human tonsil

Multiplex immunolabeling on FFPE human tonsil sections was performed by sequential IF staining on COMET™ using the FFeX technology previously described by Lunaphore Technologies^13, 20^.

## Data and code availability

The main R scripts used in this paper was deposited to Github: https://github.com/edicliuyang/Hiplex_proteome. The sequencing data reported in this paper are available upon reviewer’s request.

## Acknowledgments

We thank the Yale Center for Research Computing for guidance and use of the research computing infrastructure. The molds for microfluidic devices were fabricated at the Yale University School of Engineering and Applied Science (SEAS) Nanofabrication Center. Next-generation sequencing was conducted at the Yale Center for Genome Analysis (YCGA) as well as the Yale Stem Cell Center Genomics Core Facility which was supported by the Connecticut Regenerative Medicine Research Fund and the Li Ka Shing Foundation. Service provided by the Genomics Core of Yale Cooperative Center of Excellence in Hematology (U54DK106857) was used. This research was supported by Packard Fellowship for Science and Engineering (to R.F.), Stand-Up-to-Cancer (SU2C) Convergence 2.0 Award (to R.F.), and Yale Stem Cell Center Chen Innovation Award (to R.F.). It was also supported by grants from the U.S. National Institutes of Health (NIH) (U54AG076043 to R.F., S.H., J.E.C., and M.X.; UG3CA257393, R01CA245313, and R01MH128876, to R.F.). Y.L. was supported by the Society for ImmunoTherapy of Cancer (SITC) Fellowship.

## Contributions

Conceptualization: R.F.; Methodology: Y.L., G.S., and X.Q.; Experimental Investigation: Y.L., M.D., H.A., M.T., P.B., M.C., and M.X.; Data Analysis: Y.L., M.D., M.S., P.B., M.C., and R.F.; Resources: G.S., A.E., X.Q., and Y.D.; S.H., J.E.C, and D.H. provided valuable inputs and guidance; Original Draft: Y.L. and R.F. All authors reviewed, edited, and approved the manuscript.

## Supplemental Information

**Figure S1.**
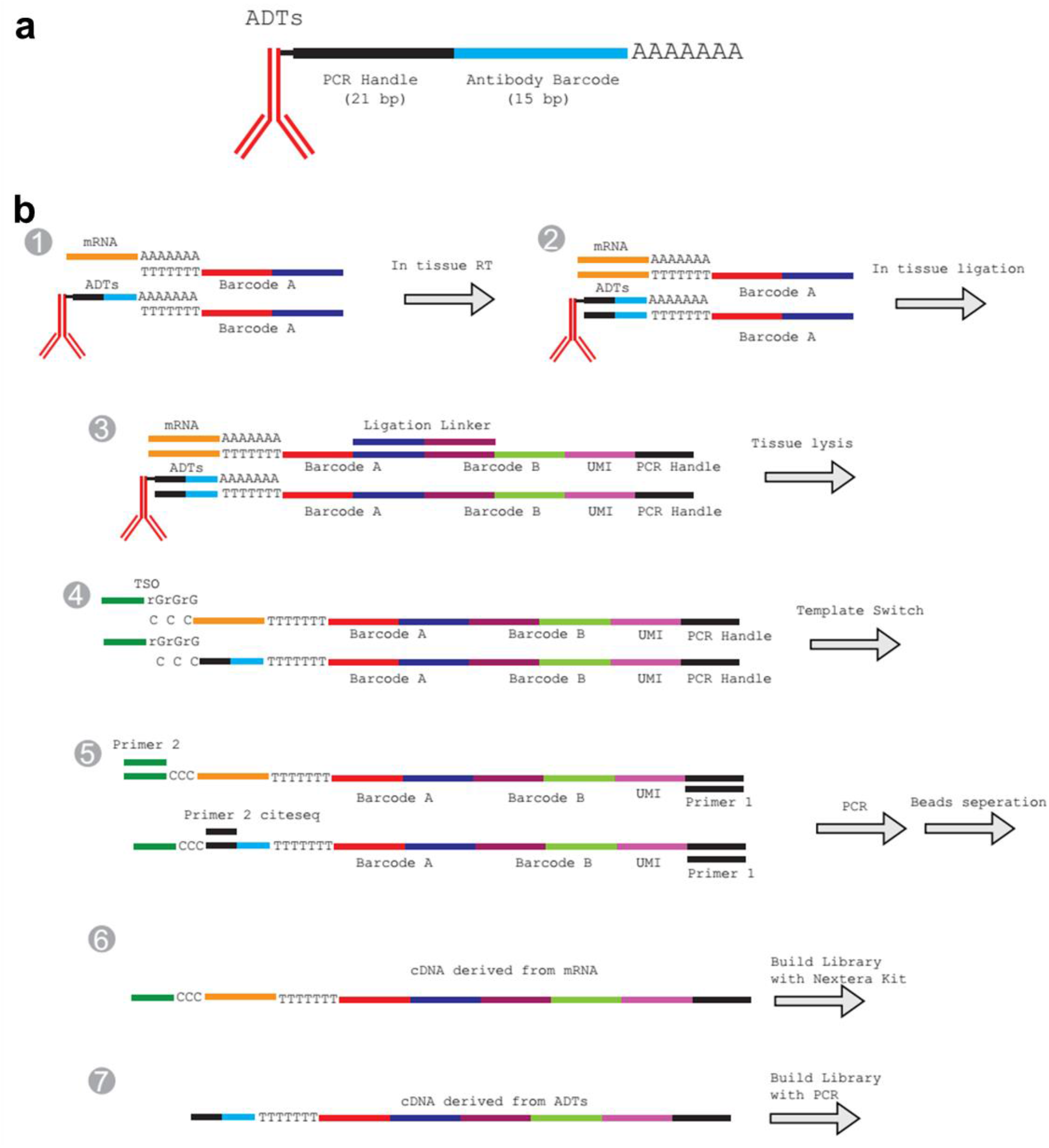
Spatial-CITE-seq design and detailed workflow. (a) ADT structure. The oligo labelled to the antibody has three functional regions: PCR handle (21 bp), antibody barcode (15 bp) and poly-A region (32 bp). (b) ADTs and mRNA with Poly-A region at the 3’ end can be reverse transcribed into cDNA using Barcode A as the RT primer. Barcode A consists of three functional regions, the poly-T region, spatial barcode region and the ligation region. During the first flow, 50 Barcode As were loaded into 50 parallel channels and the RT reaction was carried out inside each isolated channel (Step 1&2). After peeling off the 1^st^ PDMS, a 2^nd^ PDMS was attached. The in-channel ligation was performed with injecting 50 Barcode Bs into each of the 50 channels which are perpendicular to the channels of 1^st^ PDMS chip (Step 3). Barcode B has four functional regions: ligation region, barcode region, UMI region and PCR handle region. Barcode B was also 5’ biotin modification. After ligation, tissue was lysed, and cDNAs were purified with streptavidin beads. The cDNAs on the beads were templated switched with template switch oligo (Step 4). PCR was used to amplify the cDNA (Step 5). The products were split into two portions, the mRNA derived cDNAs and the ADT derived cDNAs. The library was then built separately. More details were in the method section.

**Figure S2.**
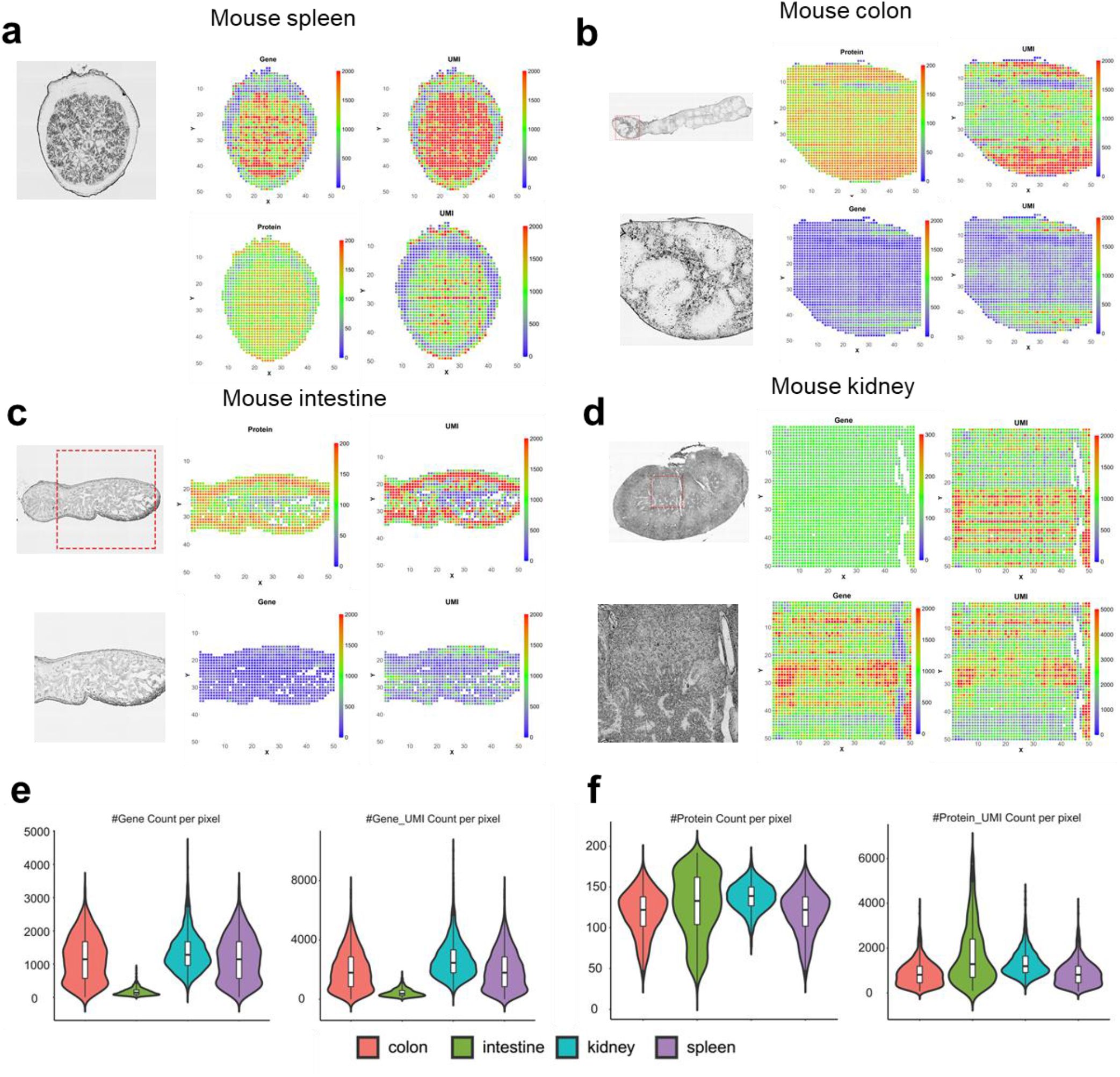
Spatial mapping of mouse spleen, colon, intestine and kidney with Spatial-CITE- seq. A 189 antibodies cocktail was used for all four mouse samples. The bright field image, spatial gene heatmap, spatial gene UMI heatmap, spatial protein heatmap and spatial protein UMI heatmap of spleen (a), colon (b), intestine (c) and kidney (d). (e) gene and gene UMI count per pixel of all four mouse samples. (f) Protein and protein UMI count per pixel of all four mouse samples.

**Figure S3.**
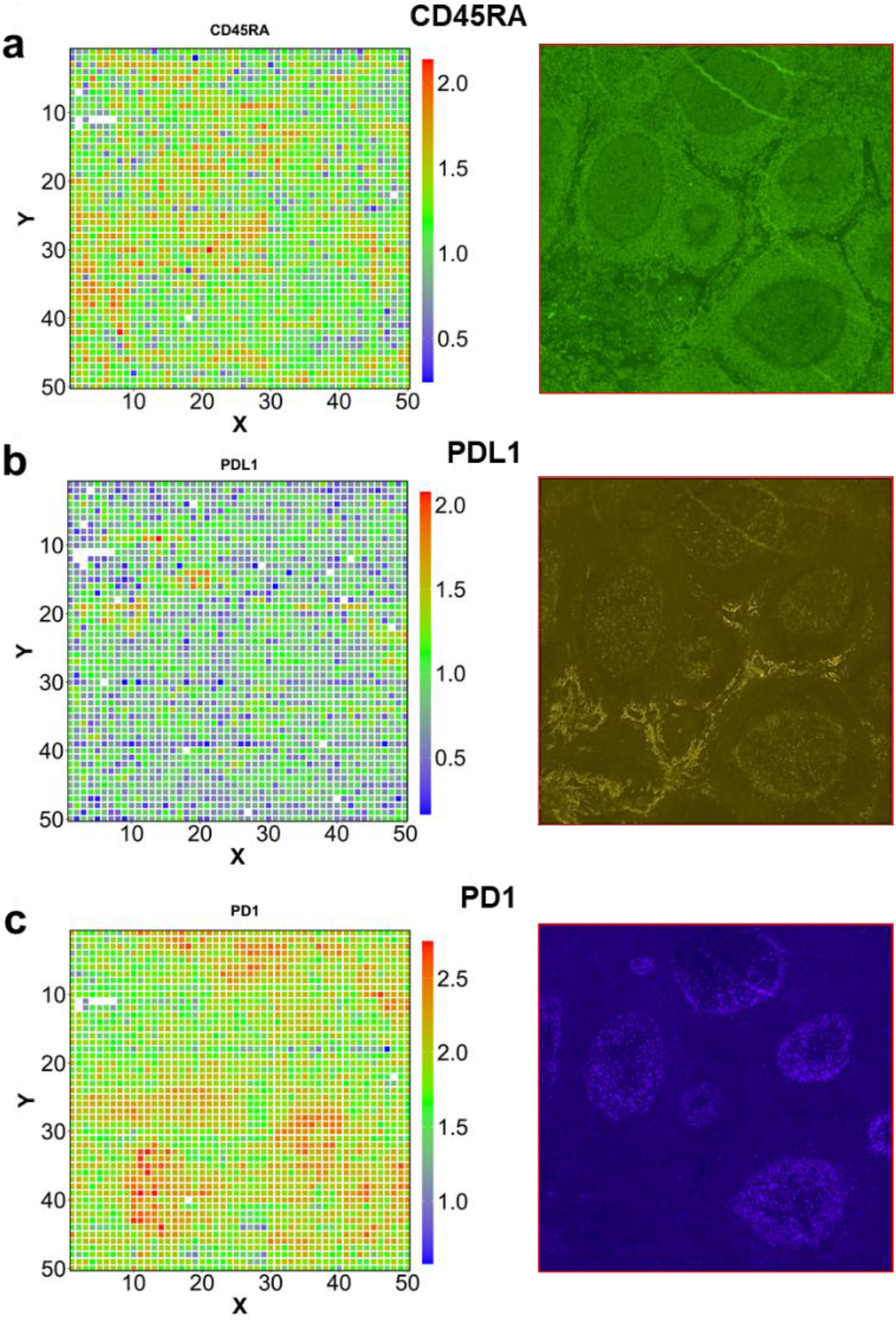
Immunostaining validation of spatial protein profiles. Sequential IF staining data using the FFeX technology by Lunaphore Technologies were compared side by side with Spatial- CITE-seq data. (a) CD45RA, (b) PDL1, (c) PD1. Note: the staining is not on exactly the same tonsil block.

**Figure S4.**
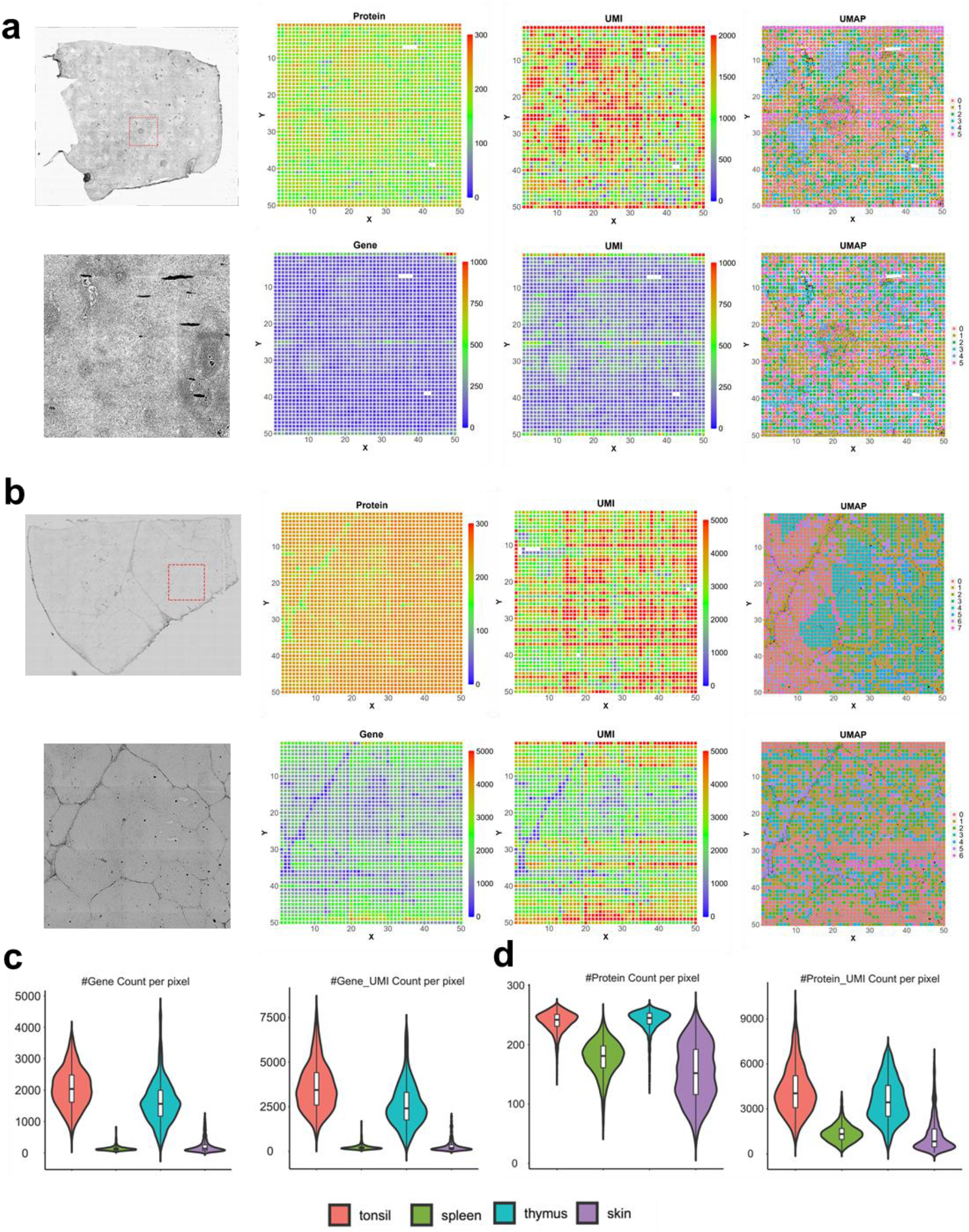
Spatial mapping of human spleen and thymus with Spatial-CITE-seq. A 273 antibodies cocktail was used for all four human samples. The bright field image, spatial gene heatmap, spatial gene UMI heatmap, spatial protein heatmap, spatial protein UMI heatmap, spatial clustering (based protein) and spatial clustering (based on RNA) of spleen (a) and thymus (b). (c) gene and gene UMI count per pixel of all four human samples. (d) Protein and protein UMI count per pixel of all four human samples.

**Figure S5.**
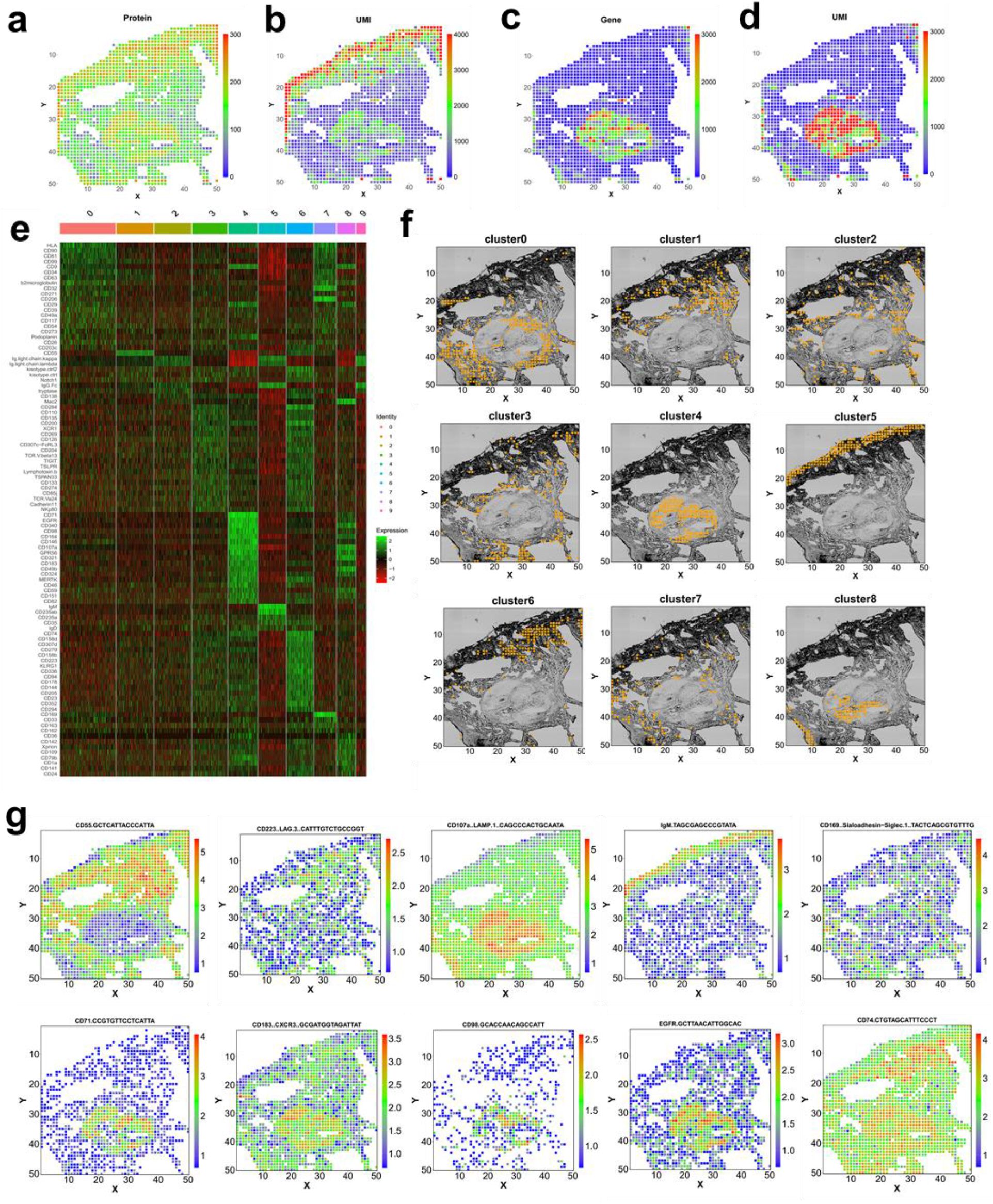
Spatial profiling of human skin biopsy tissue collected from the COVID-19 mRNA vaccine injection site. Spatial heatmap of gene (a), gene UMI (b), protein (c) and protein UMI (d). (e) Expression heatmap of the 10 clusters identified in skin biopsy sample. (f) the individual clusters plotted. (g) spatial distribution of some representative proteins.

**Figure S6.**
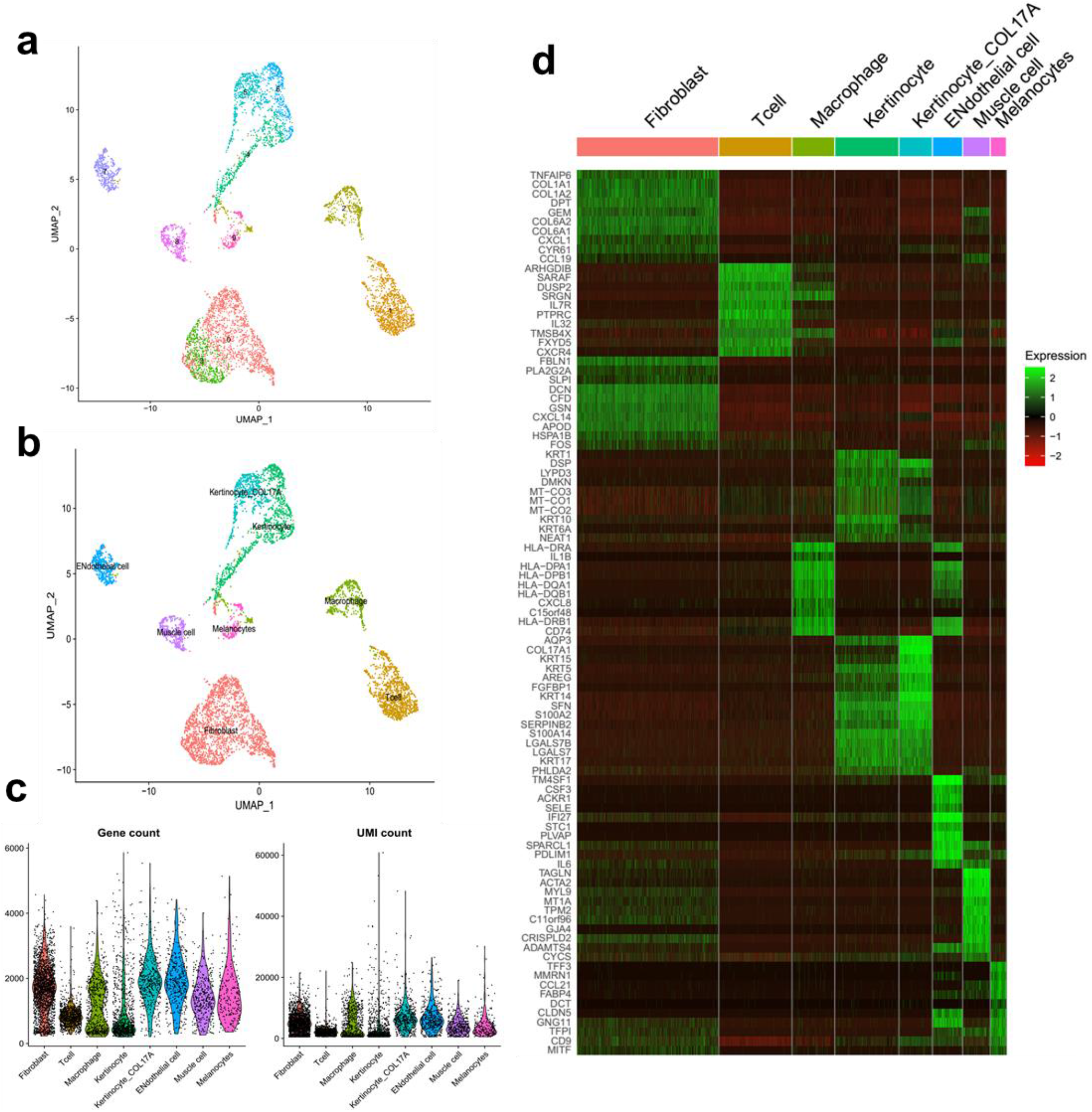
scRNA-seq sequencing data of skin biopsy sample. (a) spatial clusters of scRNA- seq data. (b) annotated cell types using canonical marker genes. (c) violin plot of genes and UMIs for each cell type. (d) Expression heatmap of different cell types.

**Table S1.** Summary of gene and protein counts for all the samples sequenced.

| Sample | # Useful Pixels | # Unique genes present | # Unique proteins present | Average # genes per pixel | Average # gene UMIs per pixel | Average # proteins per pixel | Average # protein UMIs per pixel |
| --- | --- | --- | --- | --- | --- | --- | --- |
| Spleen (Mouse) | 1303 | 19923 | 189 | 1166 | 1972 | 118 | 885 |
| Colon (Mouse) | 2037 | 19468 | 189 | 258 | 462 | 142 | 1213 |
| Intestine (Mouse) | 902 | 20444 | 189 | 172 | 447 | 129 | 1796 |
| Kidney (Mouse) | 2419 | 23750 | 189 | 1367 | 2675 | 137 | 1324 |
| Tonsil (Human) | 2492 | 28417 | 273 | 2079 | 3639 | 239 | 4309 |
| Spleen (Human) | 2494 | 20236 | 273 | 132 | 212 | 177 | 1403 |
| Thymus (Human) | 2500 | 28278 | 273 | 1647 | 2633 | 241 | 3700 |
| Skin (Human) | 1691 | 15486 | 273 | 411 | 815 | 153 | 1340 |

**Table S2.**
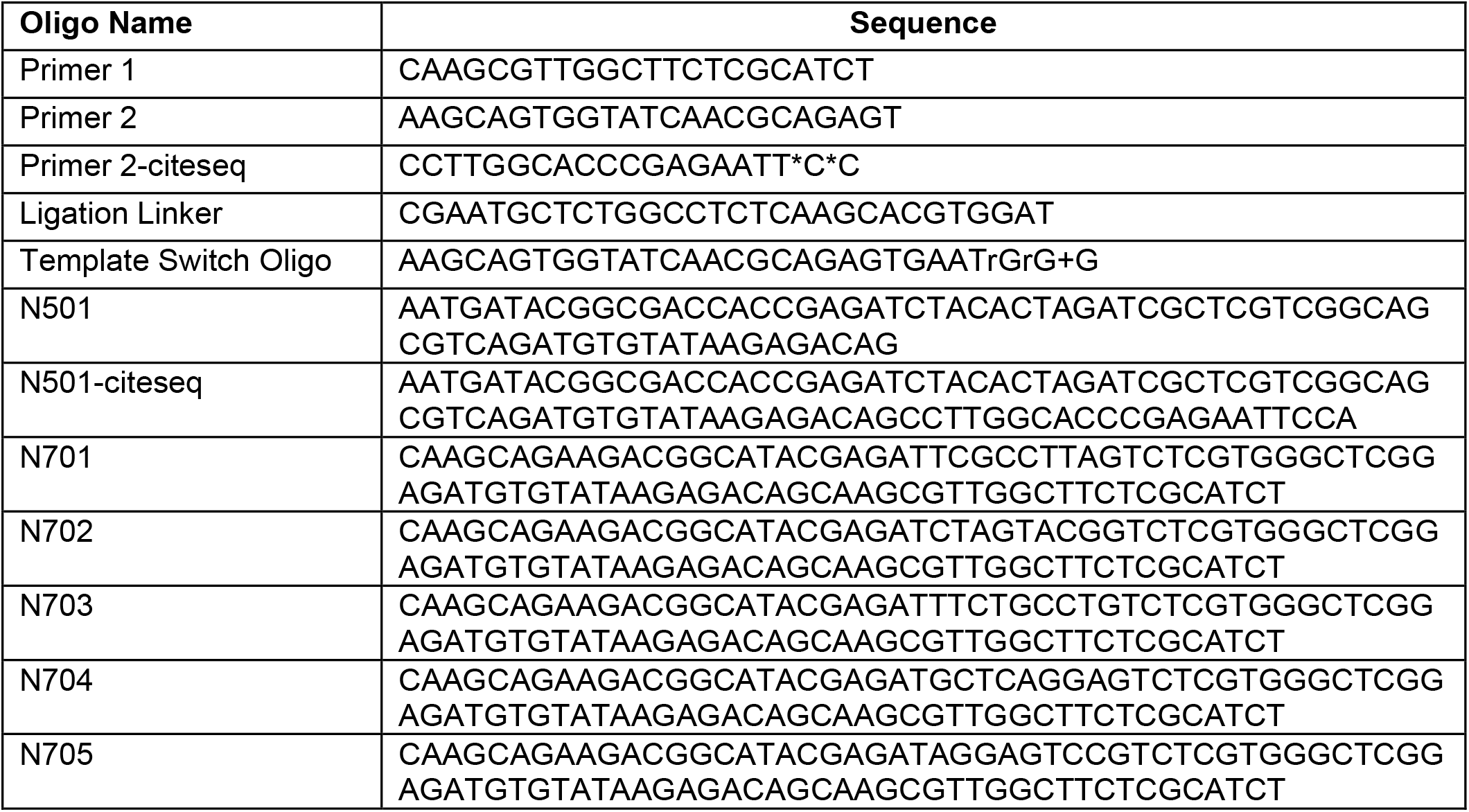
DNA oligos for PCR, ligation and library preparation. All Oligos were HPLC purified.

**Table S3.**
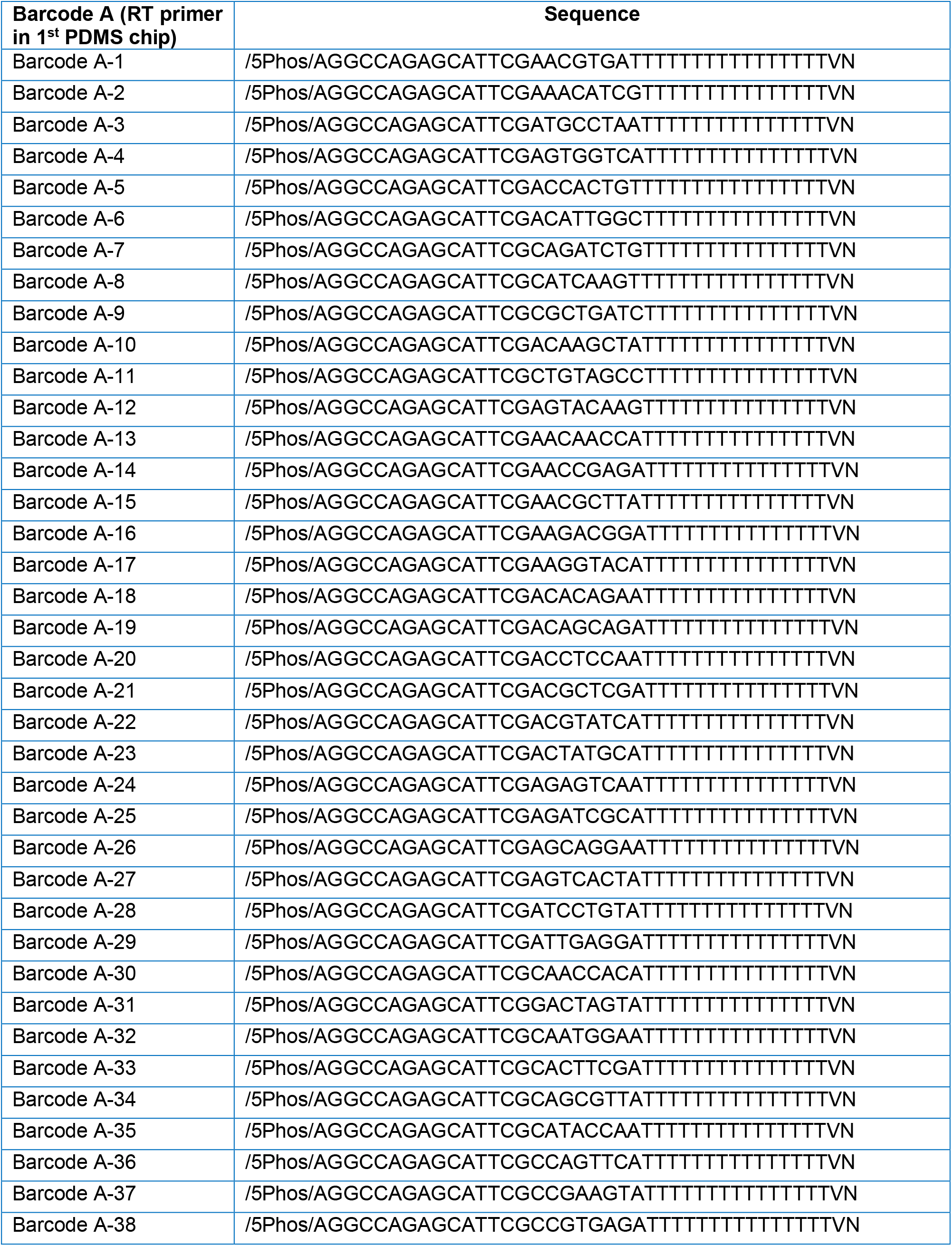
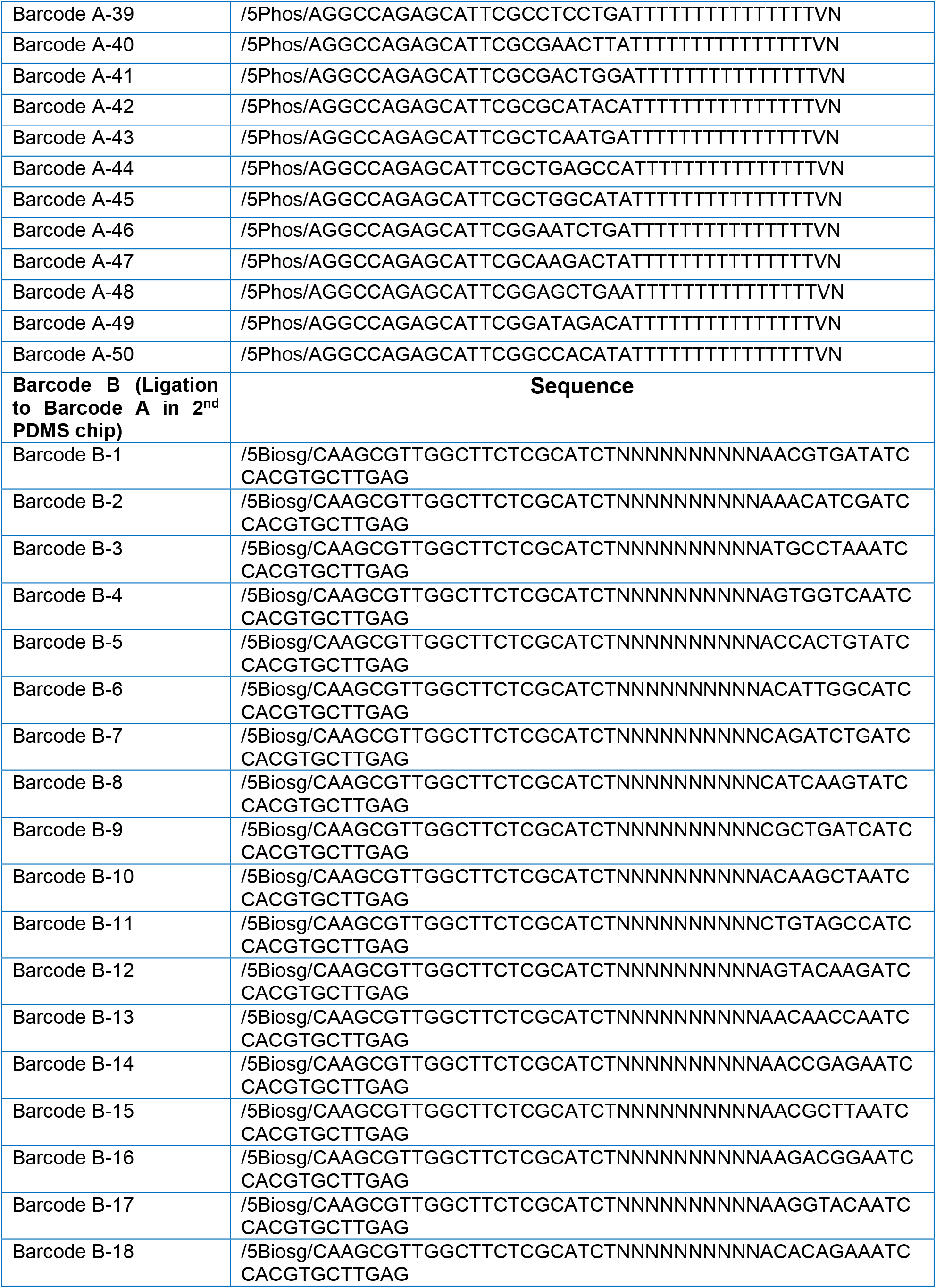
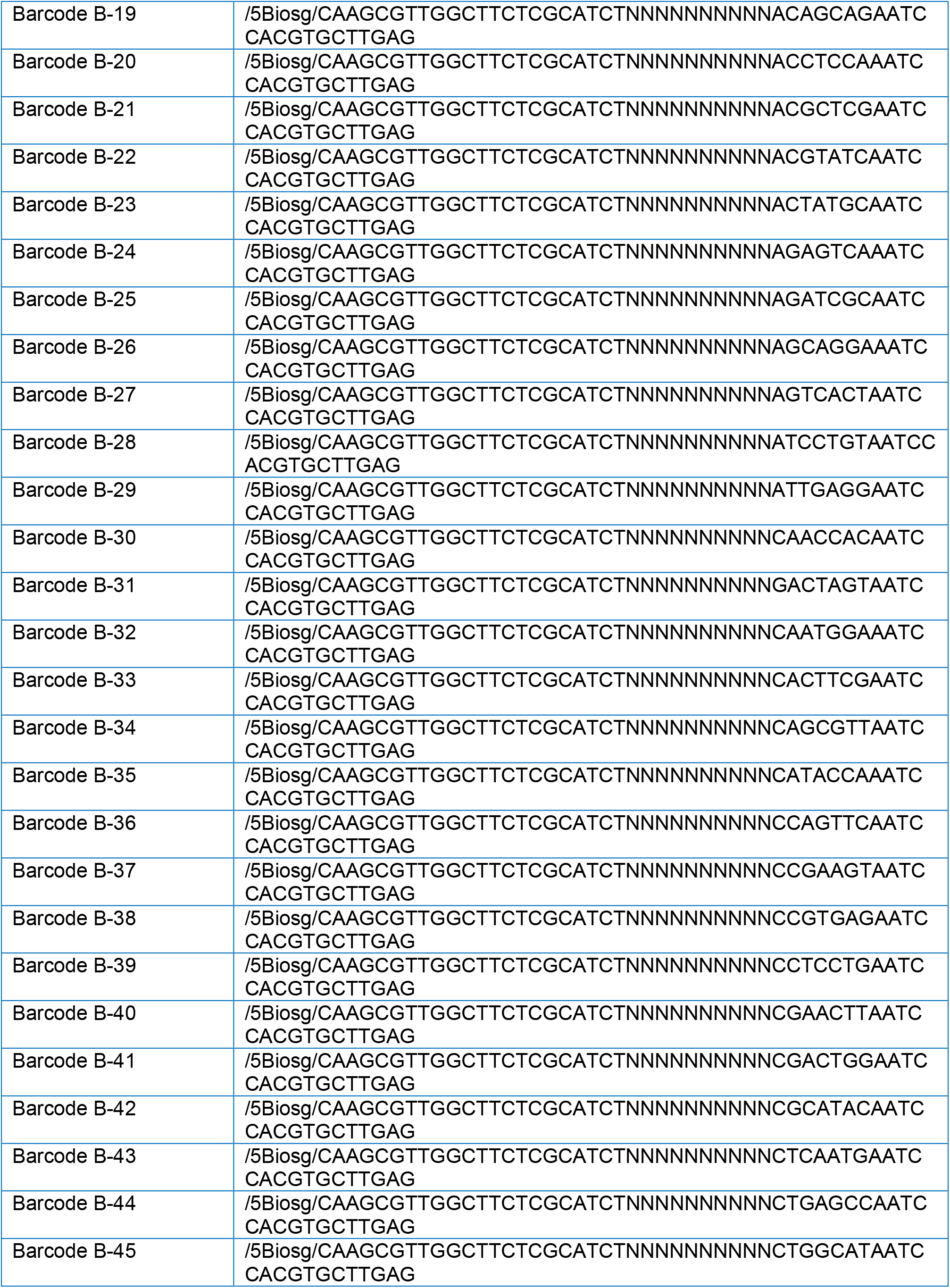
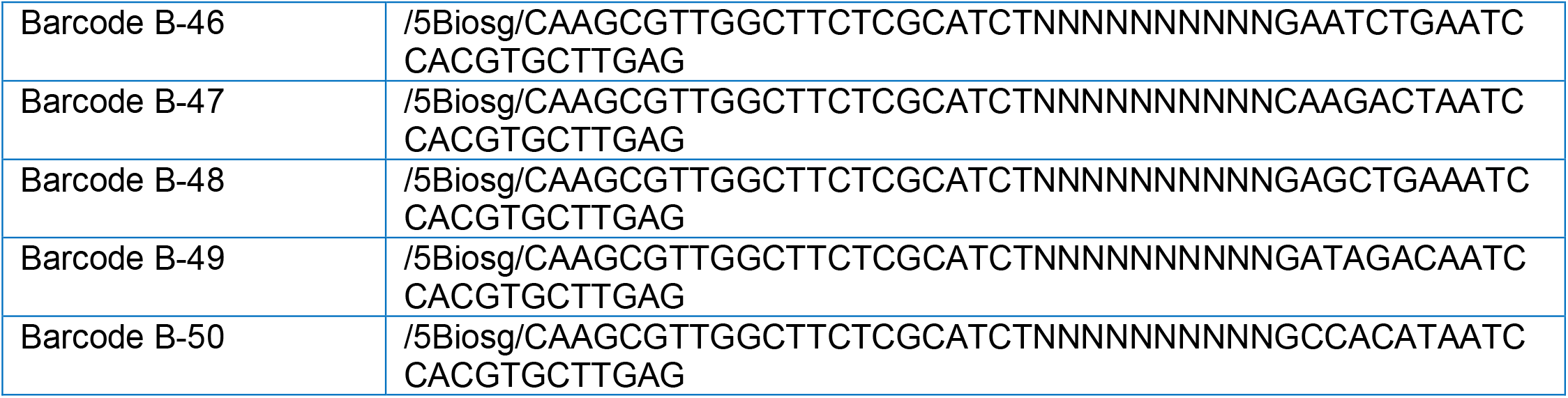
The list of 50 DNA Barcode As and 50 Barcode Bs.

**Table S4.**
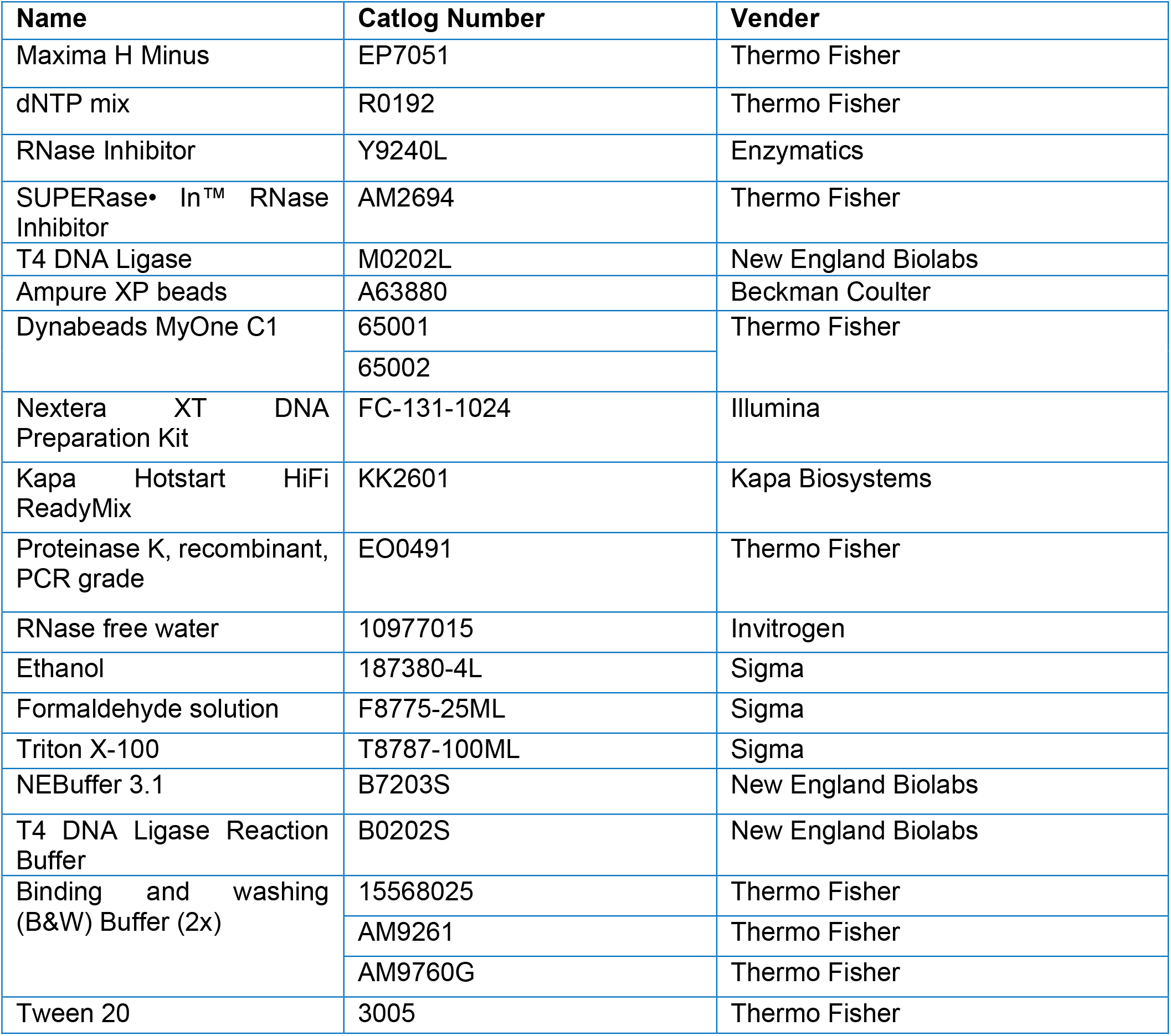
Chemicals and reagents used.

**Table S5. ADT lists for human and mouse. See attached Excel Spreadsheet - Table S5.**

## References

1. Stahl, P.L. et al. Visualization and analysis of gene expression in tissue sections by spatial transcriptomics. Science 353, 78–82 (2016).

2. Burgess, D.J. Spatial transcriptomics coming of age. Nat Rev Genet 20, 317 (2019).

3. Larsson, L., Frisen, J. & Lundeberg, J. Spatially resolved transcriptomics adds a new dimension to genomics. Nat Methods 18, 15–18 (2021).

4. Stoeckius, M. et al. Simultaneous epitope and transcriptome measurement in single cells. Nat Methods 14, 865–868 (2017).

5. Liu, Y. et al. High-Spatial-Resolution Multi-Omics Sequencing via Deterministic Barcoding in Tissue. Cell 183, 1665–1681 e1618 (2020).

6. Su, G. et al. Spatial multi-omics sequencing for fixed tissue via DBiT-seq. STAR Protoc 2, 100532 (2021).

7. Vickovic, S. et al. SM-Omics is an automated platform for high-throughput spatial multi-omics. Nat Commun 13, 795 (2022).

8. Ben-Chetrit, N. et al. Integrated protein and transcriptome high-throughput spatial profiling. bioRxiv, 2022.2003.2015.484516 (2022).

9. Carter, R.H. & Myers, R. Germinal center structure and function: lessons from CD19. Semin Immunol 20, 43–48 (2008).

10. Fischer, M.B. et al. Dependence of germinal center B cells on expression of CD21/CD35 for survival. Science 280, 582–585 (1998).

11. Santamaria, K. et al. Committed Human CD23-Negative Light-Zone Germinal Center B Cells Delineate Transcriptional Program Supporting Plasma Cell Differentiation. Front Immunol 12, 744573 (2021).

12. Takai, T. Roles of Fc receptors in autoimmunity. Nat Rev Immunol 2, 580–592 (2002).

13. Migliozzi, D. et al. Microfluidics-assisted multiplexed biomarker detection for in situ mapping of immune cells in tumor sections. Microsyst Nanoeng 5, 59 (2019).

14. Anderson, A.C., Joller, N. & Kuchroo, V.K. Lag-3, Tim-3, and TIGIT: Co-inhibitory Receptors with Specialized Functions in Immune Regulation. Immunity 44, 989–1004 (2016).

15. Yoshitomi, H. & Ueno, H. Shared and distinct roles of T peripheral helper and T follicular helper cells in human diseases. Cell Mol Immunol 18, 523–527 (2021).

16. Kuett, L. et al. Three-dimensional imaging mass cytometry for highly multiplexed molecular and cellular mapping of tissues and the tumor microenvironment. Nat Cancer (2021).

17. Goltsev, Y. et al. Deep Profiling of Mouse Splenic Architecture with CODEX Multiplexed Imaging. Cell 174, 968–981 e915 (2018).

18. Lin, J.R. et al. Highly multiplexed immunofluorescence imaging of human tissues and tumors using t-CyCIF and conventional optical microscopes. Elife 7 (2018).

19. Lin, J.R., Fallahi-Sichani, M., Chen, J.Y. & Sorger, P.K. Cyclic Immunofluorescence (CycIF), A Highly Multiplexed Method for Single-cell Imaging. Curr Protoc Chem Biol 8, 251–264 (2016).

20. Cappi, G., Dupouy, D.G., Comino, M.A. & Ciftlik, A.T. Ultra-fast and automated immunohistofluorescent multistaining using a microfluidic tissue processor. Sci Rep-Uk 9, 4489 (2019).

21. Navarro, J.F., Sjöstrand, J., Salmén, F., Lundeberg, J. & Ståhl, P.L. ST Pipeline: an automated pipeline for spatial mapping of unique transcripts. Bioinformatics 33, 2591–2593 (2017).

22. Roelli, P., bbimber, Flynn, B., santiagorevale & Gui, G. Hoohm/CITE-seq-Count: 1.4.2 (1.4.2). Zenodo (2019).

23. Stuart, T. et al. Comprehensive Integration of Single-Cell Data. Cell 177, 1888-1902.e1821 (2019).

